# Quantitative Analysis Reveals Reciprocal Regulations Underlying Recovery Dynamics of Thymocytes and Thymic Environment

**DOI:** 10.1101/478164

**Authors:** Kazumasa B. Kaneko, Ryosuke Tateishi, Takahisa Miyao, Nobuko Akiyama, Ryo Yokota, Taishin Akiyama, Tetsuya J. Kobayashi

## Abstract

Thymic crosstalk, a set of reciprocal regulations between thymocytes and thymic environment, is relevant for orchestrating the appropriate development of thymocytes as well as the recovery of the thymus from various exogenous insults. Nevertheless, the dynamic and regulatory aspects of the thymic crosstalk have not yet been clarified. In this work, we inferred the interactions shaping the thymic crosstalk and its resultant dynamics between the thymocytes and the thymic epithelial cells (TECs) by quantitative analysis and modelling of the recovery dynamics induced by irradiation. The analysis identified regulatory interactions consistent with the known molecular evidence and revealed their dynamic roles in the recovery process. Moreover, the analysis also predicted, and a subsequent experiment verified a new regulation of CD4+CD8+ double positive (DP) thymocytes, which temporarily increases their proliferation rate upon the decrease in their population size. Our model established the pivotal step towards a dynamic understanding of the thymic crosstalk as a regulatory network system.

## Main text

The thymus is an organ responsible for producing a large part of T cells with appropriate repertoires [1]. Nevertheless, it is relatively sensitive to insults by stress, virus infection, radiation, and other extra stimuli [2, 3]. While a thymus in a healthy animal can be normally recovered from these damages, a relatively prolonged process of the thymic recovery may impair T cell-mediated immunity due to a reduced replenishment of naïve T cell repertoire during the recovery period [3, 4].

Sub-lethal dose radiation on mice has been utilized as an experimental model of the thymic regeneration after insults [5, 6]. Ionizing irradiation is also broadly used for hematopoietic transplantation and cancer therapy [7, 8], and total body irradiation causes an acute thymic injury and slow recovery of thymopoiesis. Several studies showed that irradiation reduces cellularity not only of thymocytes but also of thymic epithelial cells (TECs), which are major constituents of the thymic environment [5, 9, 10]. Because the thymopoiesis is supported by interactions between the thymocytes and the TECs [11], understanding thymic recovery requires a characterization of the reciprocal regulations between the thymocytes and the TECs.

Concomitantly, various techniques to trace, perturb, and quantify the cells involved in the events have enabled us to characterize their kinetics and dynamics quantitatively [12–15]. By combining mathematical models with such quantitative data, dynamic aspects of thymopoiesis have been distilled in the forms of more detailed kinetic information, e.g., rates of proliferation, death, and differentiation [12, 16]. Mehr *et al.* [17] had developed the first kinetic model of the thymocyte development using ordinary differential equations (ODEs) [18]. Since this pioneering work, kinetic models of the thymopoiesis have been progressively refined by taking into account of the detailed cellularity and developmental states of the thymocytes and by incorporating different experimental conditions [19–23].

Nevertheless, the previous works have focused only on the thymocytes. Thymic development as well as the thymic recovery are not thymocyte-autonomous but supported by the thymic environment. In the last decade, we have accumulating molecular-biological evidences that the thymic environment itself is homeostatically maintained by the thymic crosstalk, bidirectional interactions between the thymocytes and the thymic environment [11, 24, 25]. Among several cells comprising the thymic environment, cortical and medullary thymic epithelial cells (cTECs and mTECs) play integral roles to induce and control proliferation, apoptosis, lineage commitments of thymocytes [11, 26–30]. Thymocytes, in turn, also regulate the TECs by modulating their maturation and proliferations [5, 31–33].

Despite the evident relevance and importance of the thymic crosstalk for the thymopoiesis and the thymic recovery, the kinetic aspects of the reciprocal regulations between the thymocytes and the TECs have not yet been clarified. Thus, we investigate the joint dynamics of the thymocytes and the TECs by combining a mathematical model with a quantitative measurement of the number of the thymocytes and the TECs during the recovery.

## Result

### Quantification of recovery dynamics of thymocytes and TECs

To quantitatively investigate how the thymocytes and the TECs are kinetically related and how the thymic recovery is established, we artificially perturbed the populations of the thymocytes and the TECs in thymi by using sub-lethal 4.5 Gy irradiation, and measured the dynamic changes in the population sizes of the thymocytes and the TECs after the irradiation over three weeks (Fig.1 (a)). Figure 1 (b) summarizes the changes in the numbers of cells, which were sorted based on conventional markers of the thymocytes (Fig.1 (c) and Fig. S1) and the TECs (Fig.1 (d) and Fig. S1). Figure 1 (b) shows that all types of the thymocytes and the TECs investigated decreased exponentially in number at different rates immediately after the irradiation. Then, both thymocytes and TECs started recovering at the longest within 10 days; the DN thymocytes and cTECs took only less than 5 days whereas the CD4+ SP thymocytes and the mTECs required longer intervals, which nicely reflects the temporal order of the thymocyte development from the DN to the SP cells through the interactions from the cTECs to the mTECs.

**Figure 1:**
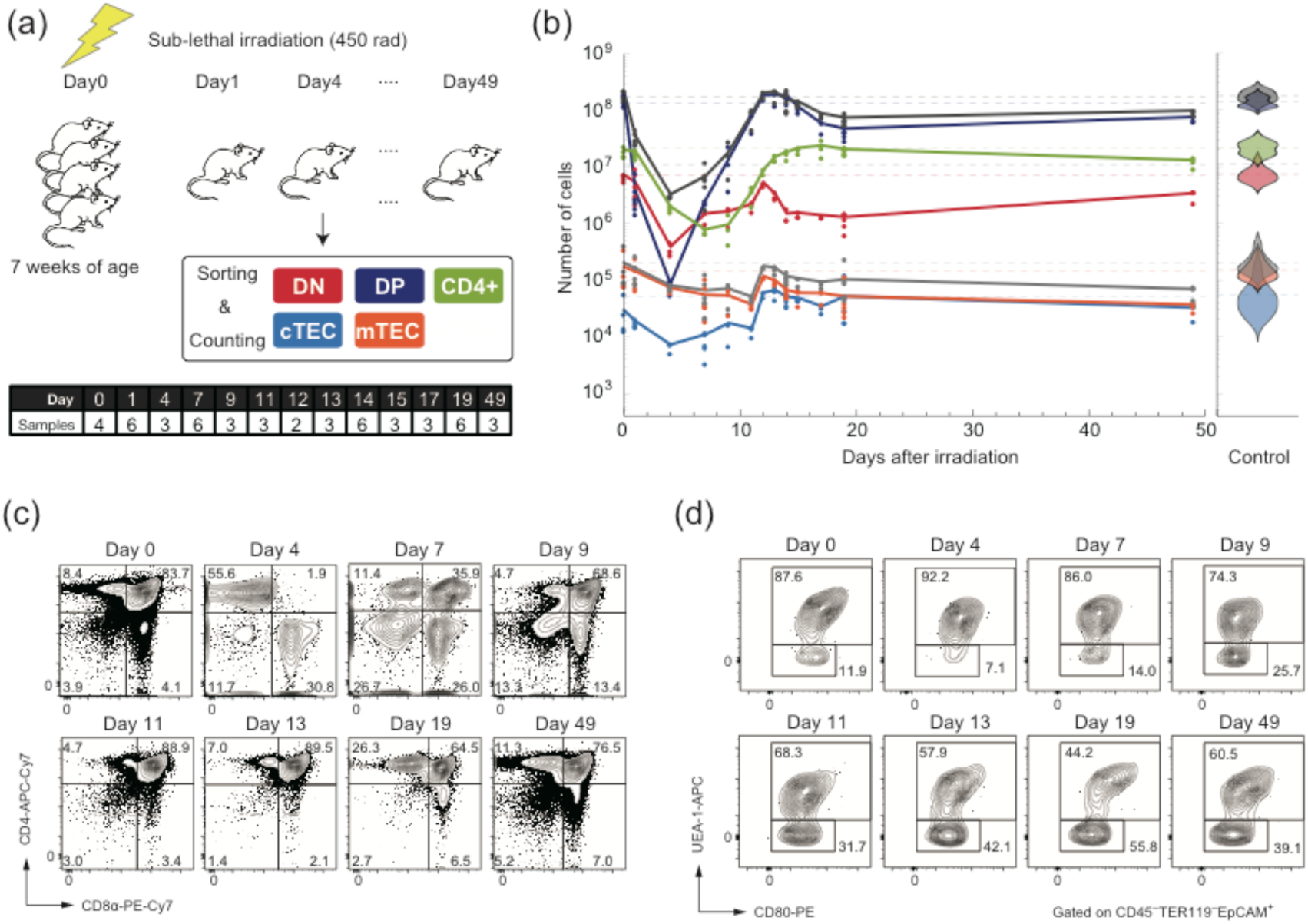
Recovery dynamics of the thymocytes and the TECs after sub-lethal irradiation. (a) A schematic diagram of the perturbation experiment and a table of the number of the sampled mice. (b) The left panel shows trajectories of the counts of the thymocytes (DN:red, DP:blue, SP4:light green) and the TECs (cTEC: cyan, mTEC: orange) after the irradiation. Points correspond to the experimental counts of the cells, and the solid curves are linear interpolations of the average counts at each time point. Dark and light gray curves represent the total numbers of thymocytes and TECs, respectively. The right panel is violin plots of the numbers of the thymocytes and the TECs without perturbation. (c) Typical flow cytometric profiles of thymocytes after the sub-lethal dose radiation. Thymocytes were analyzed by staining with anti-CD4 and anti-CD8α. Percentage of each fraction is shown in the panels. (d) Typical flow cytometric profiles of TECs after the sub-lethal dose radiation. TECs (EpCAM^+^CD45^−^ TER119^−^) were analyzed by staining with a combination of UEA-1 lectin and anti-CD80. Percentages of UEA-1^+^ cells (mTECs) and UEA-1^−^ cells (cTECs) are shown in the panels.

Upon the recoveries, the population sizes of all but the SP cells overshot around 15 days, and eventually returned to the stationary numbers, which are almost equivalent to or at least half of the original population sizes before the irradiation. Such overshooting behaviors suggest that the numbers of the thymocytes and TECs are dynamically and mutual regulated via reciprocal interactions.

### Mathematical model can reproduce the recovery dynamics

In order to infer such regulatory interactions behind the dynamics, we constructed a mathematical model for the population dynamics of the thymocytes and the TECs by ordinary differential equations (ODEs), which explicitly include five cell types: *i ∈ {DN, DP, SP4, cTEC, mTEC}*. In order to take into account of the acute influence of the irradiation to the dynamics, the number of the *i*-th cell type at time *t* (day) is decomposed into two parts; 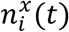 is that of the exponentially dying cells by the irradiation, and the other, *n*_i_(*t*), is that of the cells had survived from or were newly generated after the irradiation. 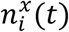 is assumed to decrease exponentially at a constant rate, *ω*_*i*_(1/day), as 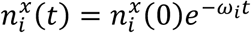, and we modelled the dynamics of *n*_i_(*t*) with ODEs. Therefore, the total number of the _i_-th cell type, 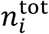, which we observed experimentally, is described as

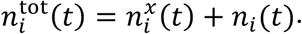

The temporal change in *n*_i_(*t*) is driven by the imbalance among influx, proliferation, death, and outflux of the *i*-th cells. While the influx can be independent of the number of the *i*-th cells, the other should, in nature, depend on the number of the existing *i*-th cells, *n*_*i*_(*t*). This allows us to generally represent the ODEs for the dynamics of *n*_*i*_(*t*) as

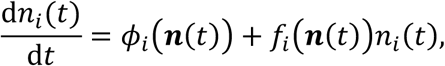

where the influx should be non-negative, *φ*_*i*_(***n***(*t*)) ≥ 0, whereas the marginalized rate of proliferation, death, and outflux *f*_*i*_(***n***(*t*)) can be either positive or negative. The actual value of *f*_*i*_(***n***(*t*)) is determined by the balance among the proliferation, the cell death, and the outflux of the *i*-th cells. In order to obtain a minimal model with a minimal complexity, we assume that both *φ*_*i*_(***n***(*t*)) and *f*_*i*_(***n***(*t*)) are at most linear with respect to ***n***(*t*) with possible constant time delays.

Therefore, our ODE model as a whole has at most quadratic nonlinearity. Even only with the quadratic nonlinearity, the fitting of the ODEs to data ends up with a non-convex and thereby hard nonlinear optimization problem, which makes an automatic parameter fitting and selection infeasible. To circumvent this difficulty, we firstly analyzed the equation only for the *i*-th cell by replacing the effects of the other cells by the actual experimental data, and narrowed down candidates of the interaction terms that appear in *φ*_*i*_(***n***(*t*)) and *f*_*i*_(***n***(*t*)) as well as their parameter values. Then, we concatenated all the candidate equations to have the following whole model:

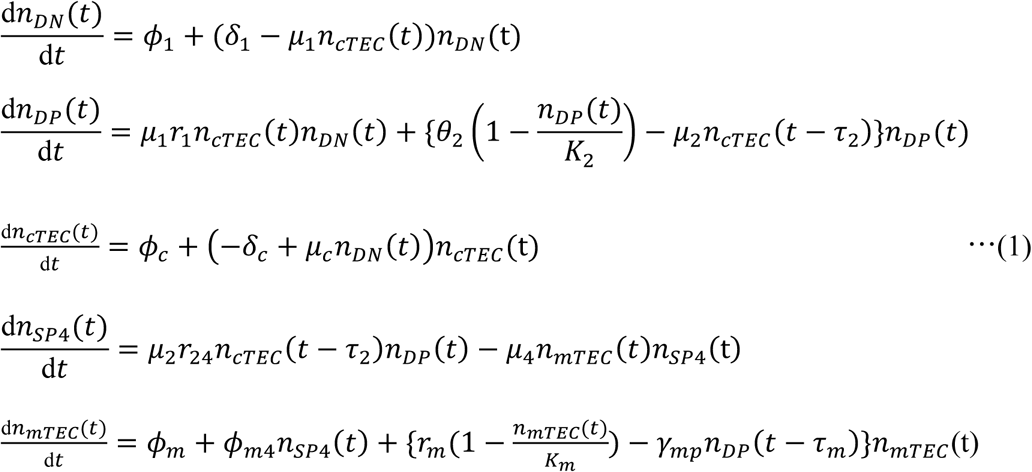

a diagrammatic representation of which is shown in Fig. 2 (a). Based on this model with the candidate parameter values as an initial condition, we conducted a nonlinear least square estimation of the parameter values so that the whole model can reproduce the experimental data (Fig. 2 (b) and Table 1). Moreover, in order to reevaluate the importance of several parameters, we statistically estimated the potential variability of the estimated values by conducting a bootstrap parameter estimation (Fig. 2 (c,d) and Table S1). As shown in Fig. 2 (b), our model, Eq.(1), nicely reproduced the recovery dynamics observed experimentally, demonstrating that the interactions depicted in Fig.2 (a) are sufficient to account for the dynamics.

**Table 1.** A comparison of the estimated kinetic rates with those from previous studies

footnote: (CI: Confidence interval) The value of each term is estimated in our model by the following equations of the parameters evaluated at the steady state $n_i^*$ , :
| Term | Value | CI | Previous Study |
| --- | --- | --- | --- |
| Inflow rate to DN [Cells/Day] | $3.30 \times 10^4$ | $[3.0 \times 10^3, 5.9 \times 10^4]$ | $2.0 \times 10^4$ [20] $1.3 \times 10^4$ [26] |
| DN apparent proliferation rate [/Day] | $1.50 \times 10^{-1}$ | $[-2.3 \times 10^{-2}, 2.9 \times 10^{-1}]$ | $2.3 \times 10^{-1}$ [20] $6.2 \times 10^{-4}$ [26]<br>$3.6 \times 10^{-1}$ [21] |
| DN differentiation rate [/Day] | $1.60 \times 10^{-1}$ | $[1.7 \times 10^{-5}, 3.0 \times 10^{-1}]$ | $2.4 \times 10^{-1}$ [20] $2.8 \times 10^{-2}$ [26]<br>$3.4 \times 10^{-1}$ [21] $4.5 \times 10^{-1}$ [27] |
| DN residence time [Hour] | $1.50 \times 10^2$ | $[8.0 \times 10^1, 1.4 \times 10^6]$ | $4.2 \times 10^2$ [20] $3.5 \times 10^2$ [21] |
| DP apparent proliferation rate [/Day] | $1.00 \times 10^{-1}$ | $[8.9 \times 10^{-2}, 8.4 \times 10^{-1}]$ | $1.5 \times 10^{-2}$ [20] $-3.7 \times 10^{-1}$ [22]<br>$-1.6 \times 10^{-1}$ [27] $-9.0 \times 10^{-3}$ [26] |
| DP differentiation rate to SP4 [/Day] | $1.10 \times 10^{-1}$ | $[9.1 \times 10^{-2}, 8.4 \times 10^{-1}]$ | $2.1 \times 10^{-2}$ [20] $1.2 \times 10^{-2}$ [22]<br>$3.0 \times 10^{-2}$ [27] $9.9 \times 10^{-2}$ [26] |
| DP residence time [Hour] | $2.20 \times 10^2$ | $[2.8 \times 10^1, 2.6 \times 10^2]$ | $9.4 \times 10^1$ [20] $7.6 \times 10^1$ [22] |
| Ratio of DP to SP4 in DP export [%] | 11 | [9.6, 75] | 6.0 [20], 0.016[22], 8.1[27],<br>65 [26] |
| SP4 apparent export rate [/Day] | $5.30 \times 10^{-1}$ | $[4.9 \times 10^{-1}, 1.5 \times 10^1]$ | $2.0 \times 10^{-2}$ [20] $9.0 \times 10^{-2}$ [22]<br>$1.4 \times 10^{-1}$ [27] $1.7 \times 10^{-1}$ [26] |

Inflow rate to DN thymocytes: $\phi_1$
DN apparent proliferation rate : $\delta_1 - \mu_1(1 - r_1)n_{cTEC}^*$
DN differentiation rate : $\mu_1 r_1 n_{cTEC}^*$
DN residence time : $24/(\mu_1 r_1 n_{cTEC}^*)$
DP apparent proliferation rate : $\theta_2(1 - n_{DP}^*/K_2) - (1 - r_{24})\mu_2 n_{cTEC}^*$
DP differentiation rate to SP4 : $r_{24}\mu_2 n_{cTEC}^*$
DP residence time : $24/(r_{24}\mu_2 n_{cTEC}^*)$
Ratio of DP to SP4 in DP export : $r_{24}$
SP4 apparent export rate : $\mu_4 n_{mTEC}^*$

**Figure 2:**
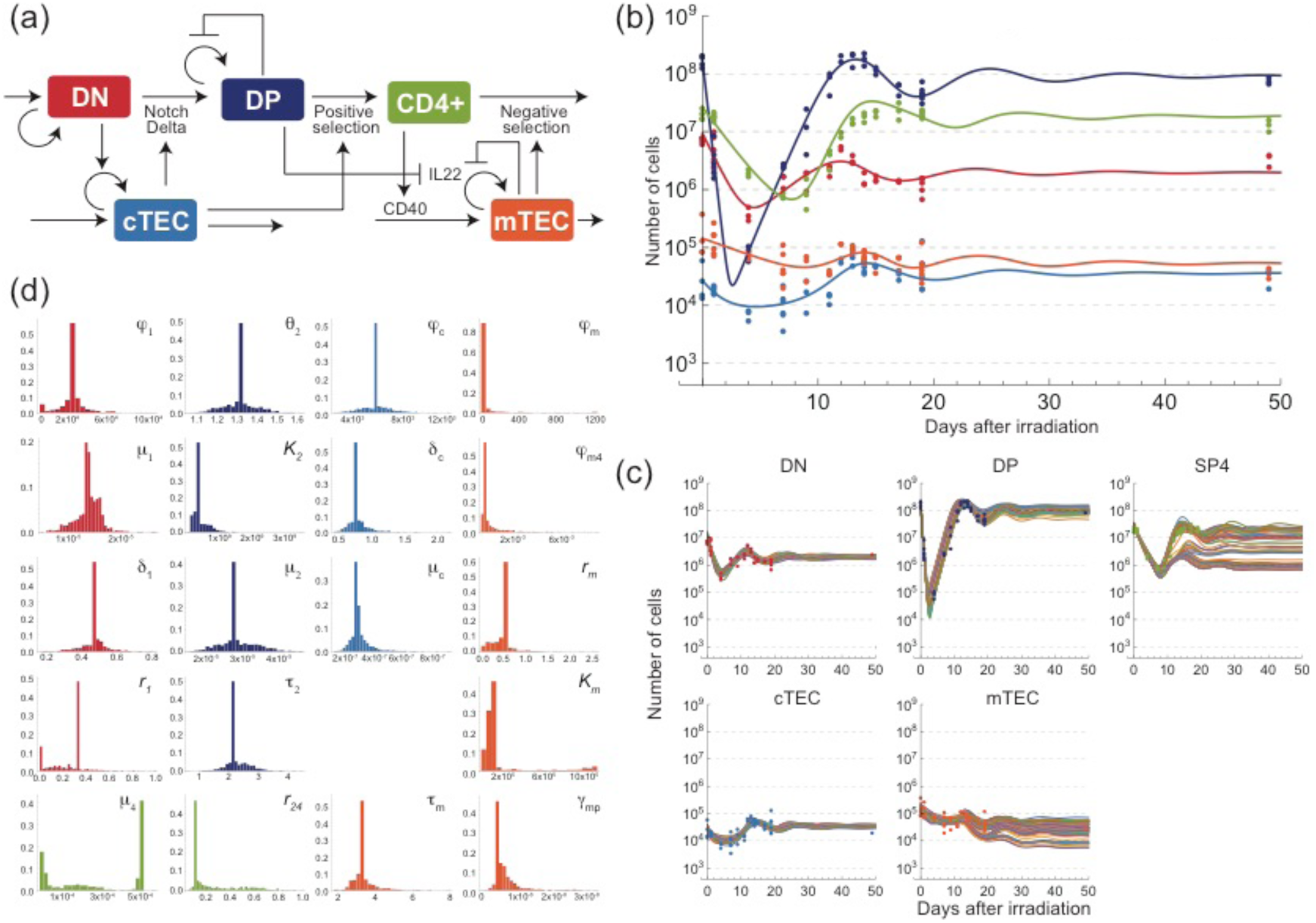
A schematic diagram, trajectories, and estimated parameter distributions of the mathematical model inferred from the quantitative data. (a) A schematic diagram of the inter cellular interactions inferred from the experimental data and implemented in Eq. (1). (b) Trajectories of the numbers of the thymocytes and the TECs obtained by simulating Eq. (1) with the optimized parameter set. The curves represent simulated trajectories, and the points are the same experimental data as in Fig. 1 (b). (c) Trajectories obtained in the bootstrap parameter estimation. The trajectories in different panels with the same color correspond to the trajectories obtained from the optimal parameter estimated from bootstrapped samples. The trajectories of 100 randomly selected samples are shown in the panels. (d) The variations of parameters estimated by the bootstrap parameter estimation. The colors of histograms designate the related cell types to the parameters. The variations of other parameters are shown in Fig. S2.

### DN thymocytes and cTECs form a negative feedback

Our estimated model indicates that the DN thymocytes and the cTECs form a negative feedback, in which the DN cells positively regulate the cTECs whereas the cTECs effectively inhibit the increase in the DN cells (Fig. 2 (a)). This negative feedback is the source of the overshooting behaviors in the recovery dynamics, and can account for the lag in the onset of the cTECs recovery by few days behind that of the DN cells.

These interactions inferred from the quantitative recovery data are also consistent with molecular evidences identified previously. On the one hand, the positive interaction from the DN thymocytes to the cTECs may be interpreted as the induction of the cTEC proliferation by the DN cells, which was evidenced by the fact that the number of the mature cTECs decreases if the DN differentiation is blocked at early stages [31, 34]. On the other hand, our model suggests that the cTECs down-regulate the number of the DN cells. This negative regulation is a marginal effect of induced cell death, induced differentiation from the DN to the DP stages, and inhibition of DN proliferation by the cTECs. This negative regulation of the DN cells by the cTECs is quite consistent with the Notch1-Delta-like4-dependent lineage commitment of the DN cells to the DP stage mediated by the cTECs [27, 28]. It should be noted however that our model does not exclude other possibilities of additional molecular interactions as long as their marginal influences are consistent with the diagram in Fig. 2 (a).

### DP recovery is achieved by temporal up-regulation of proliferation

The kinetic component characteristic to the DP dynamics is its much faster recovery than that of the DN cells (Fig. 1 (b)), which strongly suggest that the DP recovery is achieved by a self-proliferation rather than the influx from the DN population. However, we have inconsistent evidences on the self-proliferation ability of the DP cells and its speed, which might depend on strains [35]; some of which showed that the DP cells proliferate little [22, 36] whereas others suggest the DP cells can proliferate faster than the other types of the thymocytes [35, 37]. Our model coordinates these properties by an auto-inhibitory regulation of the DP proliferation represented by the logistic term in Eq.1, which can realize a fast proliferation during the recovery period and its slowdown at the steady state. Nevertheless, such auto-inhibitory regulation in the DP proliferation has not yet been reported.

To experimentally verify this prediction by our model, we estimated the fraction of proliferating DP cells under the same condition as in Fig. 1(a) by staining the DP cells with a proliferation marker Ki67 (Fig. 3(a)). We observed that the fraction of the proliferating DP cells transiently increased and peaked at Days 7 after the irradiation, which perfectly coincides with the timing of the exponential increase in the DP cells during the recovery. The self-proliferation ceased as the number of the DP cells had recovered to the normal population size before the irradiation. This result strongly supports that the number of the DP cells is negatively regulated by its total population size to maintain its homeostasis. In addition, this autoregulatory mechanism is consistent with the previous observations that the DP cells proliferate little when they are at the steady state number [36].

**Figure 3:**
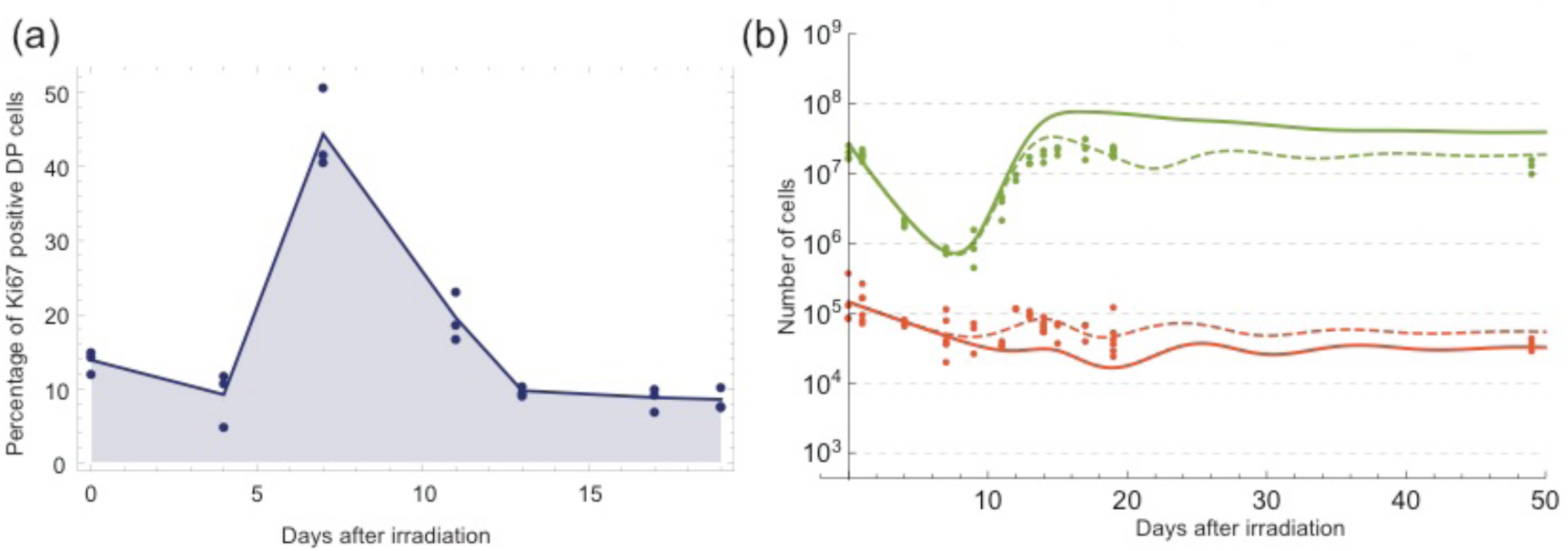
Validations of the model prediction by proliferation assay of DP cells and *in silico* evaluation of impact from the disturbed crosstalk between SP4 thymocytes and mTECs. (a) Percentages of Ki67 positive DP cells 0, 4, 11, 13,17, and 19 days after the irradiation. Points are experimental counts of the cells, and the shaded lines are linear interpolations of the average counts. (b) Simulated trajectories of the SP4 thymocytes and the mTECs (Thick solid curves) with the parameter values mimicking the experimental condition in [33], γ_4_ = 8.0 × 10^−6^ and *φ*_m4_ = 0. The thin dashing curves are those obtained with the optimal parameter values used in Fig. 2 (b).

While the autoregulatory proliferation of the DP cells is necessary for reproducing the fast recovery, it alone cannot account for the overshooting behavior of the DP cells, which suggests other regulations of the DP cells by other cells. Supported by the well-established evidences that the cTECs engage in the positive selection of the DP cells, our model includes a negative regulation of the DP cells by the cTECs with a time-delay, which can nicely reproduce the overshoot of the DP cell count. This negative regulation can be interpreted as the marginal effect of an induced apoptosis of the DP cells with non-functional TCRs and the differentiation of the DP cells into the single positive cells upon the rescue of the apoptosis.

Our model estimates that the fraction of the rescued DP cells that differentiated into CD4 SP, *r*_24_, is about 11%, which is within the range of the previous estimates that 0.02~65% of DP cells survive and differentiate into CD4 SP via positive selection (Table 1). We should note that the estimated ratio of the rescued DP cells varies in the previous studies; mostly because the apoptosis rate cannot be estimated directly only from the dynamics of the population sizes, including our case. To partially circumvent this problem, we also estimate that the stable rate of the DP cells to differentiate into CD4 SP, 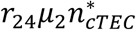 is 1.10×10^−1^ (1/day), which is closer to the range of the previous estimates from 1.2×10^−2^ to 9.9×10^−2^ (1/day) (Table 1).

### DP and CD4 SP thymocytes incoherently regulate mTEC recovery

Compared with the other thymocytes and the TECs, the CD4 SP cells showed a much slower recovery with a less pronounced overshooting (Fig. 1(b)). This slow recovery of the CD4 SP cells is consistent with the lack of a proliferation capacity of the CD4 SP cells [13, 35], which leads to a prolonged recovery. The CD4 SP dynamics can be reproduced by no proliferation and mTEC-dependent death and outflux that represent the negative selection of the SP cells by the mTECs (Fig. 2(a)).

In contrast, the mTEC recovery was initiated almost concurrently with that of the cTECs (Fig. 1(b)). While interactions with the CD4 SP cells have been proven to be essential for the maturation of the mTECs [38], the prolonged CD4 SP recovery is not sufficient for reproducing the earlier onset of the mTEC recovery. Our model incorporates an auto-inhibitory regulation of the mTEC proliferation and its negative regulation by the DP cells with a time delay as in Eq. (1). The auto-inhibitory regulation is necessary because we obtained biologically inconsistent parameter values in the mTEC dynamics without the regulation (Fig. 4 (a) (e)). The negative regulation by the DP cells is also responsible for the overshoot of the mTECs. These mechanisms are supported by preceding experimental investigations [5, 39]. The percentage of the Ki67hi mTECs was shown to increase only after the depletion of the mTECs in [39], which suggests the existence of the auto-inhibitory regulation. In [5], the DP cells were suggested to negatively regulate the TECs proliferation via an IL22 dependent manner by using a depletion experiment of the DP cells. The DP-dependent regulation was not the only interaction that could explain the early onset of the mTEC recovery. We had also found that a DN-dependent regulation could reproduce it (Fig. 4 (b) (f)). However, this possibility was excluded in our model because we lack molecular evidence that can support the long-range interaction from the DN cells to the mTECs, which reside in spatially segregated areas of a thymus.

**Figure 4:**
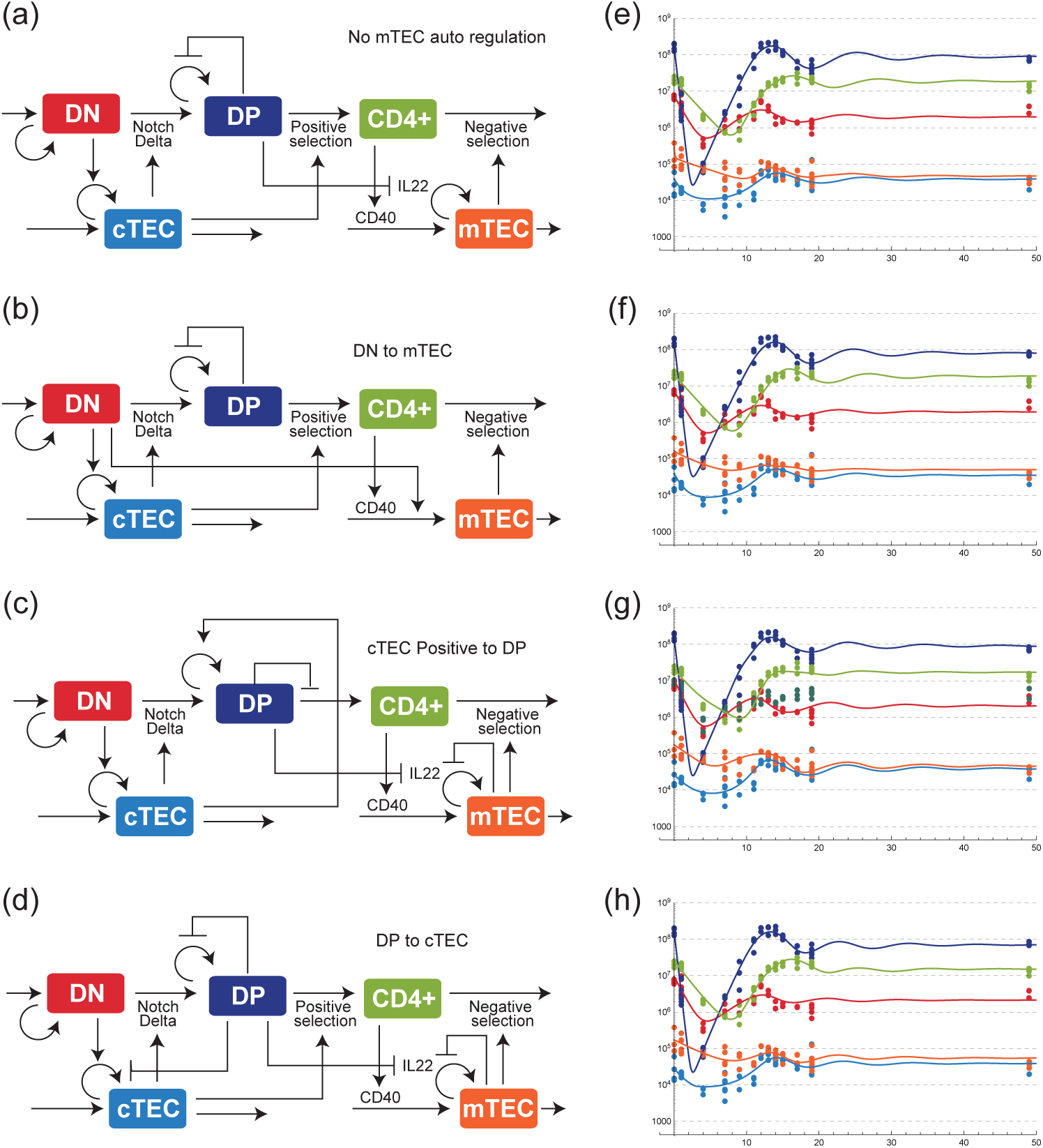
Possible regulatory mechanisms that can also reproduce the recovery dynamics of the data, but are biologically less relevant than our proposed model in Fig. 2(a). Each model incorporates the following different interactions from the model in Fig. 2(a): (a) This model excludes autoinhibitory regulation of mTEC, whereas the model in (b) includes an upregulation from DN to mTEC instead of the downregulation from DP in Fig. 2(a). The model in (c) assumes an upregulation from cTEC to DP rather than the downregulation, and that in (d) includes an upregulation from DP to cTEC. (e), (f), (g), and (h) show the corresponding trajectories of the models in (a), (b), (c), and (d), respectively.

Along with the regulated proliferation, our model assumes reciprocal regulations between the mTECs and the CD4 SP cells in order to account for the evidences that the mTEC maturation is also related with the CD4 SP cells. According to [33], the mTECs express a ligand CD80 and CD86 and a receptor CD40, the corresponding receptor and ligand of which are respectively CD28 and CD40L mainly in the CD4 SP T cells. A knock-out of CD80, CD86 and CD40 was shown to decrease the number of the mTECs and double the number of the CD4 SP cells. We substituted smaller values of *μ*_4_ and *φ*_m4_ than the estimated values to reproduce the experiment in [33] by assuming that the KO of CD80, CD86 and CD40 corresponds to this substitution. The result qualitatively reproduced the KO mutant result in [33]; the stable number of CD4 SP cells became twice whereas the number of the mTECs became about half as in Fig. 3 (b).

## Discussion

From a quantitative time-series data of the thymocyte and the TEC recoveries after a X-ray perturbation, we constructed a mathematical model for the recovery dynamics of the thymocytes and the TECs. The model fairly well reproduces the transient dynamics of the population sizes of the cells, and most of the interactions identified by the modeling are consistent with known molecular evidences.

Since previous works on quantitative characterizations of the thymocyte development using mathematical models focused only on the dynamics of the thymocyte, our work, which additionally includes both the dynamics of and the interactions with the TECs, can be viewed as an extended model of those works [17, 19–21, 40, 41]. We validated that the estimated parameter values of our model are mostly consistent with those estimated in the previous works (Table 1). Few mismatches of the parameter values might be attributed to the differences in the experimental setting and conditions.

The thymic crosstalk includes various signaling pathways, which indicates the complex regulations behind the population size control of the thymocytes and the TECs. Because of this complexity, our model may contain missing interactions or other possibility of different regulations, part of which were tested in the process of the model identification. For example, the cTECs rescue the DP thymocytes from the apoptosis via the positive selection, which leads to the increase in the DP population size. Concurrently, the positive selection also induces the differentiation of the DP cells to the SP stage, which decreases the DP population size. These contradictory interactions introduce the possibility that the cTECs upregulate the DP thymocytes rather than inhibition as being assumed in our model. We examined the possible upregulations on the DP thymocytes by the cTECs and concluded that the downregulation, which our model assumed, is more valid because of the upregulation resulted in much higher parameter values than expected from the previous works (Fig. 4 (c) (g)). We also investigated a model in which the DP thymocytes contribute to the recovery not only of the mTECs but also of the cTECs [5]. We found that the estimated parameter for the interaction from the DP cells to the cTECs was almost 0 (Fig. 4 (d) (h)), which does not support a major contribution of the DP cells to the cTEC recovery under our experimental condition.

Our model can also provide explanations of the mechanisms how specific dynamics appear in the recovery dynamics and their potential biological functions; the overshoots of the DN thymocytes and the cTECs may originate from the negative feedback between them and may contribute to a prompt recovery from various perturbations affecting the numbers of the thymocytes and the TECs. Similarly, the disinhibition of the DP proliferation upon the decrease in the DP population size facilitates the swift recovery of the DP cells, which could not be achieved only by the influx of the DN cells, the population size of which is much smaller than that of the DP cells. Our model provides an integrative view on the thymic crosstalk as a regulatory network, and serves as a starting point for a comprehensive understanding on homeostasis of thymic development.

Nevertheless, our model still has rooms for future improvement by accommodating the more detailed information on the cellularity of the thymic resident cells, such as B cells, dendritic cells, and thymic endothelial cells. These cells may have different roles in the dynamic regulation of thymic homeostasis than thymocytes and TECs, although we did not explicitly include them by presuming that their effects to the number of the thymocytes or the TECs are relatively small or constant, which was implicitly modeled by the constant parameters in our model. Actually, BMP4 production by the endothelial cells after irradiation, which can contribute to the recovery of the TECs, was reported to be constant when normalized by the size of thymus [42]. An explicit incorporation of these cells can be crucial for extending our model to other experimental setting than ours and also for deriving a more integrative and comprehensive model of the thymic development and homeostasis. Among others, the repertoires of the thymocytes are of particular relevance. The TECs are not only responsible for controlling the number of the thymocytes but also for selecting the thymocytes with appropriate repertoires. Upcoming challenges may be an integrative modeling and analysis of the thymic homeostasis in both the number and the repertoire of the cells by combining quantitative measurement and high-throughput sequencing [43].

## Methods

### Ethics statement

Animals used in the present study were maintained in accordance with the “Guiding Principles for Care and Use of Animals in the Field of Physiological Science” set by the Physiological Society of Japan. All animal experiments were approved by the Animal Research Committees of RIKEN.

### Mice, X-ray irradiation, and Flow Cytometory

Balb/cA mice were purchased from CLEA Japan. Mice (7 weeks-old) received X-ray radiation (4.5 Gy). At each sampling point after the irradiation, the mice were sacrificed, and their thymi were used for a flow cytometric analysis. Each thymus was cut and gently agitated in 2 ml of RPMI-1640 (Sigma-Aldrich, St. Louis, MO, U.S.A.) to release thymocytes for the flow cytometric analysis. The days of measurement and the number of the sampled mice are shown in Fig. 1 (a). The rest of the thymic tissue was digested using Liberase in RPMI1640 (Wako) at 37 degree for 30 min. The thymic stroma rich-fraction was analyzed by flow cytometry to detect the TEC populations. For the flow cytometric staining, cells were pre-treated with anti-CD16 and CD32 (Biolegend) for 20 min and subsequently stained with fluorescence-labeled antibodies in phosphate buffered saline containing 3% fetal bovine serum. The stained cells were analyzed by Canto II (BD). The total thymic cell numbers were evaluated by adding cell numbers of the thymic stroma-rich fraction and the thymocyte fraction. Since the DN population contains other minor populations of cells such as dendritic cells, the cell numbers of these fractions were subtracted from the DN cell number in the mathematical modeling based on the average percentage of these cells (16.6%) in the DN fraction under a steady condition. The TECs were defined as CD45-TER119-EpCAM+ cells. The mTECs and the cTECs were separated with UEA-1 staining. PECy7-anti-CD4, APCCy7-anti-CD8, APCCy7-anti-CD45, APCCy7-anti-TER119, FITC-anti-EpCAM, PE-anti-CD80 and Streptavidin-PECy7 were purchased from Biolegnd. UEA-biotin was from Vector laboratories (Burlingame, CA).

### Estimation of proliferating DP cells

Thymocytes were pre-treated with anti-CD16 and CD32 (Biolegend) and subsequently stained with anti-CD4 and anti-CD8 antibodies in phosphate buffered saline containing 3% FBS. The cells were fixed and permeabilized with Foxp3/Transcription Factor Staining Buffer Set (eBioscience) according to the manufacturer’s protocol. After the fixation and the permeabilization, the cells were stained with a PE-labeled anti-Ki67 antibody (Biolegend) and subsequently analyzed by Canto II (BD).

### Mathematical Modeling of Thymocyte and TEC dynamics

We assume that the total number of the type *i* cells, 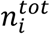, is the sum of dying cells by the irradiation 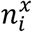 and survived or newly generated cells *n*_*i*_:

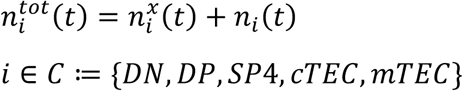

where *C* is defined as the set of the cell types.

We describe the decrease in the irradiated cells by an exponential decay, which assumes that cells die at a constant rate *ω*_*i*_ after the irradiation:

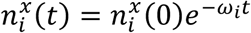

In the modeling, 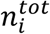 represents the initial population size of the *i*th cells and *p*_i_ is assumed to be the fraction of the survived cells at *t* ≤ 0 as

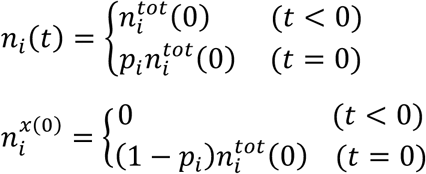

Given these initial conditions, the model of Eq (1) was implemented on MATLAB (R2016b; The MathWorks, Natick, MA) and numerically simulated by ‘dde23’ function or on Mathematica (version 11.2; Wolfram research, Champaign, Illinois) and simulated by ‘NDSolve’ function.

### Parameter estimation

In the parameter estimation, all the parameters that appear in Eq (1) and *ω*_*i*_, 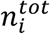, and *p*_*i*_ were estimated simultaneously. The parameters were estimated by minimizing the sum of the squares of difference between the logarithm of the observed data and the simulated values of the model. More specifically, for the observed time points *t*^∗^ = [*t*_1_, …, *t*_*m*_] and the corresponding data points *N*_*i*_(*t*_*j*_) for all *i* ∈ *C*, the estimated parameter set 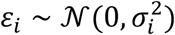 was obtained by solving

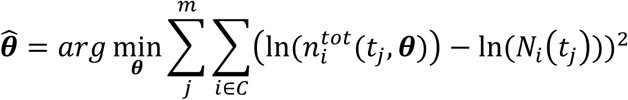

To solve this minimization, we used ‘lsqnolin’ function in MATLAB Optimization Toolbox in which the parameters were estimated by Trust Region Reflective method. The initial values of the parameters in the estimation were given so that the result converges to moderate values considering the results of related previous works. The searching range of each parameter except *p*_*i*_, *r*_1_ and *r*_24_ was set between 100 times and 0.01 times of the initial value. Since *p*_*i*_, *r*_1_ and *r*_24_ represent fractions, their searching ranges were set between 0 and 1. The symbols, the descriptions, and the estimated values of the parameters are listed in Table. S1.

### Confidence Interval by bootstrap

We calculated the confidence interval of the estimated parameter values by a bootstrap method [44].

First, for the *i*th type of the cells, we modeled the statistical variation of the data points by a Gaussian random variable 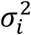 with mean 0 and variance 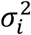 as

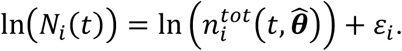

We estimated 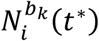 by the sample variance as

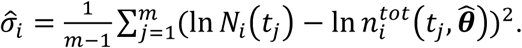

We obtained the *k*th bootstrapped sample 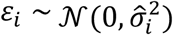 by using a random number 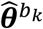 as

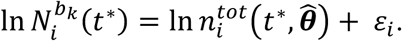

The *k*th bootstrapped parameter set 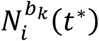 was obtained by solving the same optimization problem of the previous section by replacing the data with the *k*th bootstrapped sample 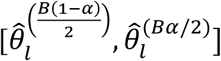 as

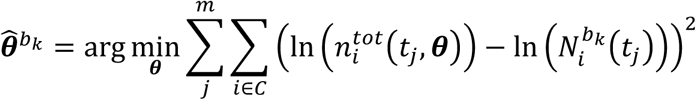

The total number of the bootstrapped samples generated was *B* = 1000. The two sided *α* ∗ 100% confidence interval of the *l*th parameter was calculated as 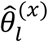 where 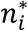 is the *x*th smallest value of the *l*th parameter obtained from the bootstrapped samples. The confidence interval of each parameter is shown in Table S1. A pairwise scatter plot of the bootstrap estimated values is shown in Fig. S3. The trajectories of the cells obtained from 100 samples of the bootstrap parameter sets are shown in Fig. 2 (c).

## Supporting information

## Data and code availability

All data and codes used in this paper are provided upon a request to the authors.

## Acknowledgements

We acknowledge Yuki Sughiyama, Masashi K. Kajita for fruitful discussions. This research was supported by Grant-in-Aid for Exploratory Research (15K14433), Grant-in-Aid for Scientific Research B (17KT0014) from the Ministry of Education, Culture, Sports, Science, and Technology, Japan, JST PRESTO Grant Number JPMJPR15E4, Japan, a grant from The Uehara Memorial Foundation, Grant-in-Aid for Scientific Research on Innovative Area from MEXT (18H04989) and Grant-in-Aid for Scientific Research from JSPS (17H04038).

## Author Contributions

T. Akiyama and T. J. Kobayashi designed the study. R. Tateishi performed experiments. T. Miyao, N. Akiyama assisted in experiments. K. Kaneko and T. J. Kobayashi performed mathematical modeling and analyzed data, and R. Yokota provided critical comments on modeling and analysis. K. Kaneko, T. Akiyama, and T. J. Kobayashi wrote the manuscript. K. Kaneko and R. Tateishi equally contributed to this work.

