## Supplementary material for "Quantitative Analysis Reveals Reciprocal Regulations Underlying Recovery Dynamics of Thymocytes and Thymic Environment"

### Supplemental Information

### SI Figures

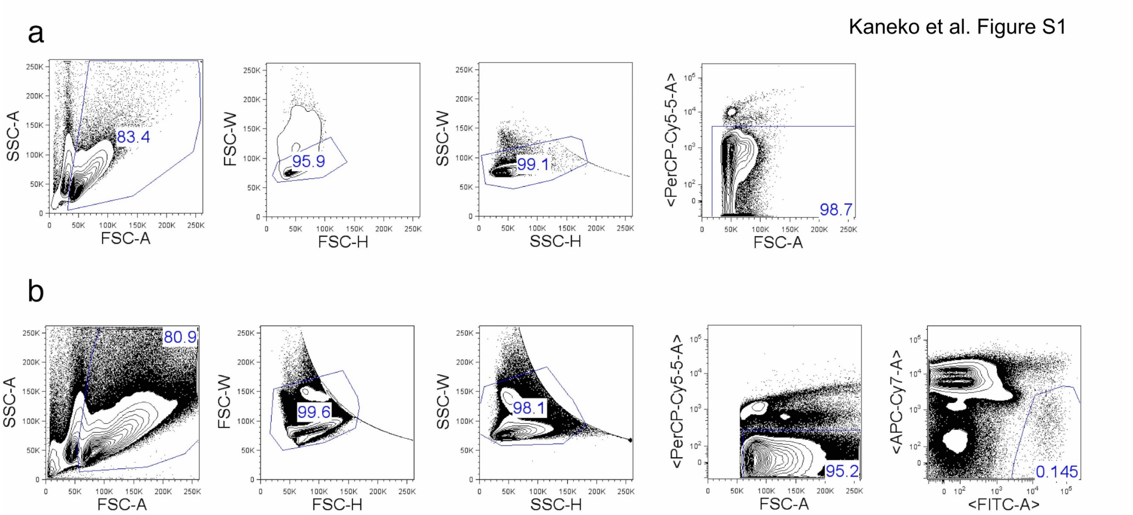


Figure S1: (a) Gating strategy for thymocytes. (b) Gating strategy for TECs.

Kaneko et al. Figure S2

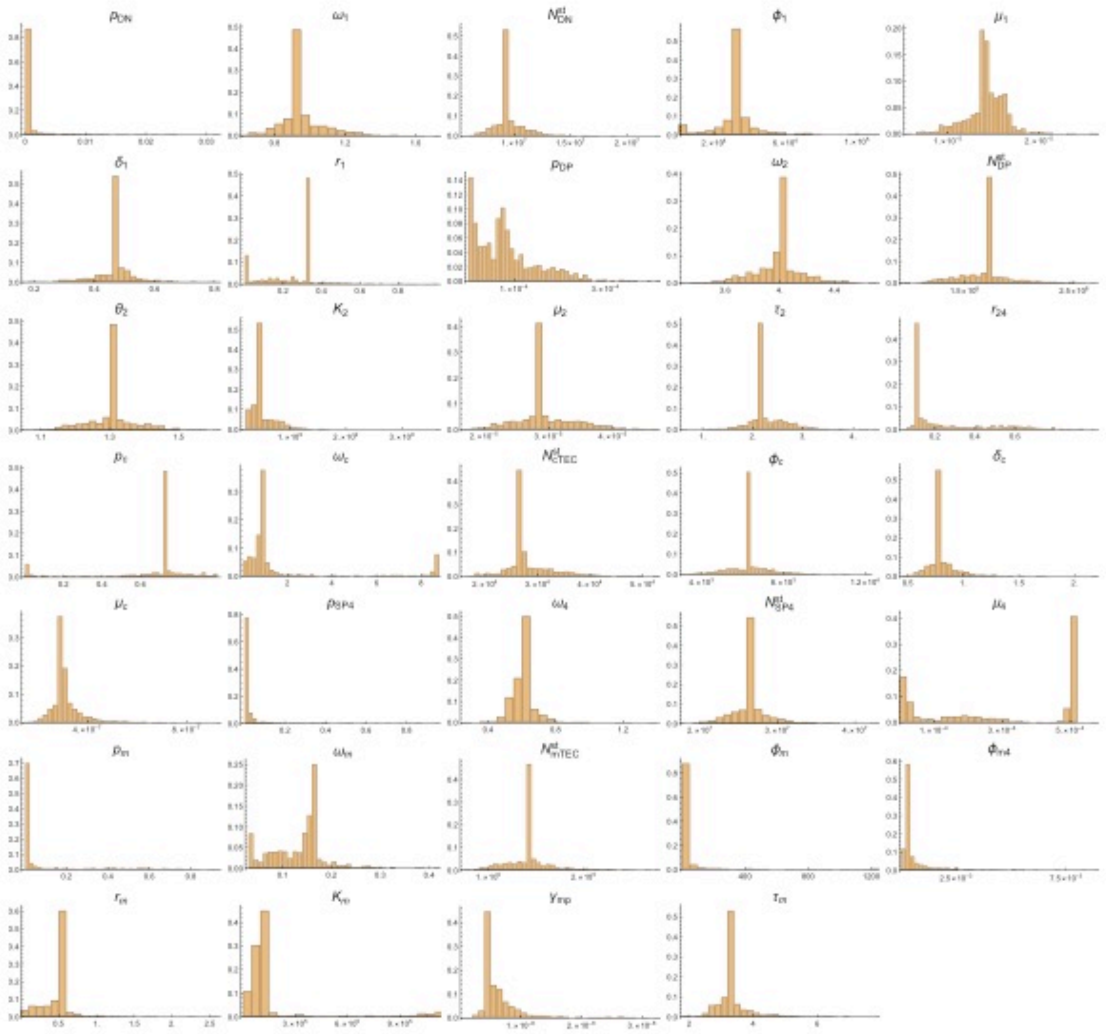

Figure S2: Histograms of the parameter values obtained by the bootstrap estimation.

Kaneko et al. Figure S3

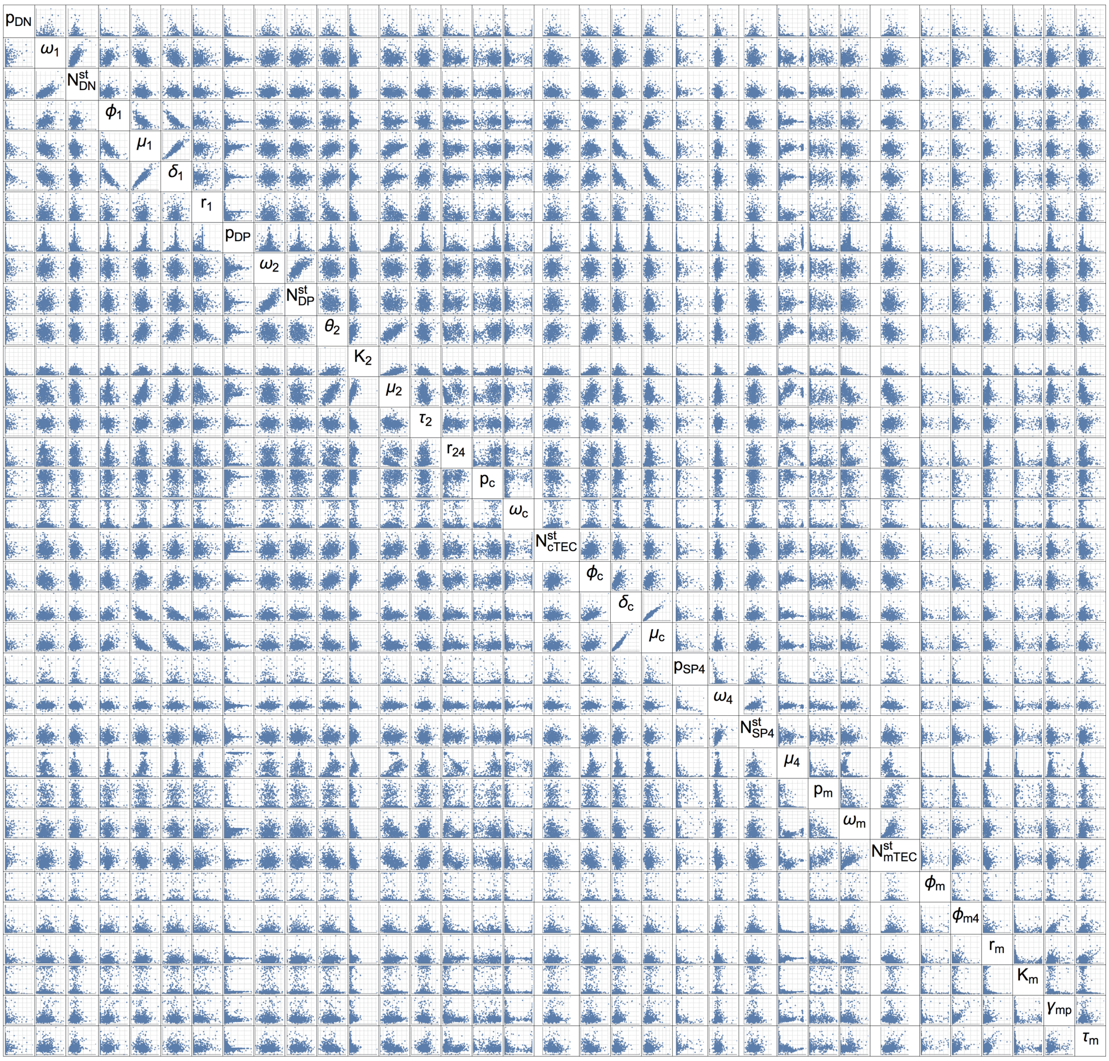


Figure S3: Two dimensional scatter plots of the parameter values obtained by the bootstrap estimation. The plot range of each parameter is set to be the same as that in Fig. S2.

### SI Tables

Table S1. Estimated parameters in the model. (CI : Confidence interval)

| Symbol | Description | Value | CI |
|---|---|---|---|
| $p_{DN}$ | Ratio of normal cells in the initial number of DN | $2.62 \times 10^{-6}$ | $[6.90 \times 10^{-7}, 9.14 \times 10^{-3}]$ |
| $\omega_{DN}$ | Death rate of irradiated DN | $9.24 \times 10^{-1}$ | $[7.35 \times 10^{-1}, 1.28 \times 10^{0}]$ |
| $n_{DN}^{tot}(0)$ | Initial value for the number of DN | $9.29 \times 10^{6}$ | $[7.15 \times 10^{6}, 1.29 \times 10^{7}]$ |
| $\phi_1$ | Inflow to DN | $3.30 \times 10^{4}$ | $[3.00 \times 10^{3}, 5.89 \times 10^{4}]$ |
| $\mu_1$ | Negative regulation by cTEC to DN | $1.38 \times 10^{-5}$ | $[9.17 \times 10^{-6}, 1.84 \times 10^{-5}]$ |
| $\delta_1$ | Intrinsic proliferation rate of DN | $4.76 \times 10^{-1}$ | $[3.00 \times 10^{-1}, 6.17 \times 10^{-1}]$ |
| $r_1$ | Ratio of differentiation to DP in regulation by cTEC | $3.33 \times 10^{-1}$ | $[4.24 \times 10^{-5}, 5.75 \times 10^{-1}]$ |
| $p_{DP}$ | Ratio of normal cells in the initial number of DP | $3.09 \times 10^{-6}$ | $[1.04 \times 10^{-7}, 2.48 \times 10^{-4}]$ |
| $\omega_{DP}$ | Death rate of irradiated DP | $4.01 \times 10^{0}$ | $[3.59 \times 10^{0}, 4.40 \times 10^{0}]$ |
| $n_{DP}^{tot}(0)$ | Initial value for the number of DP | $1.71 \times 10^{8}$ | $[1.21 \times 10^{8}, 2.28 \times 10^{8}]$ |
| $\theta_2$ | Proliferation rate of DP | $1.30 \times 10^{0}$ | $[1.15 \times 10^{0}, 1.46 \times 10^{0}]$ |
| $K_2$ | Carrying capacity of DP | $4.10 \times 10^{8}$ | $[2.42 \times 10^{8}, 1.04 \times 10^{9}]$ |
| $\mu_2$ | Negative regulation rate by cTEC to DP | $2.85 \times 10^{-5}$ | $[2.10 \times 10^{-5}, 3.97 \times 10^{-5}]$ |
| $\tau_2$ | Delay of cTEC regulating DP | $2.15 \times 10^{0}$ | $[1.60 \times 10^{0}, 3.16 \times 10^{0}]$ |
| $r_{24}$ | Ratio of differentiation to CD4+SP in regulation by cTEC | $1.06 \times 10^{-1}$ | $[9.56 \times 10^{-2}, 7.53 \times 10^{-1}]$ |

| | | | |
|---|---|---|---|
| $p_{cTEC}$ | Ratio of normal cells in the initial number of cTEC | $7.33 \times 10^{-1}$ | $[1.36 \times 10^{-3}, 9.58 \times 10^{-1}]$ |
| $\omega_{cTEC}$ | Death rate of irradiated cTEC | $8.14 \times 10^{-1}$ | $[1.02 \times 10^{-1}, 8.70 \times 10^{0}]$ |
| $n_{cTEC}^{tot}(0)$ | Initial value for the number of cTEC | $2.70 \times 10^{4}$ | $[2.05 \times 10^{4}, 4.04 \times 10^{4}]$ |
| $\phi_{c}$ | Inflow to cTEC | $6.26 \times 10^{3}$ | $[4.19 \times 10^{3}, 8.61 \times 10^{3}]$ |
| $\delta_{c}$ | Death rate of cTEC | $7.62 \times 10^{-1}$ | $[5.87 \times 10^{-1}, 1.19 \times 10^{0}]$ |
| $\mu_{1}$ | Positive regulation by DN to cTEC | $3.00 \times 10^{-7}$ | $[2.18 \times 10^{-7}, 5.25 \times 10^{-7}]$ |
| $p_{SP4}$ | Ratio of normal cells in the initial number of CD4+SP | $1.40 \times 10^{-2}$ | $[1.83 \times 10^{-4}, 3.50 \times 10^{-1}]$ |
| $\omega_{SP4}$ | Death rate of irradiated CD4+SP | $6.23 \times 10^{-1}$ | $[4.91 \times 10^{-1}, 7.72 \times 10^{-1}]$ |
| $n_{SP4}^{tot}(0)$ | Initial value for the number of CD4+SP | $2.64 \times 10^{7}$ | $[2.16 \times 10^{7}, 3.27 \times 10^{7}]$ |
| $\mu_{4}$ | Negative regulation by mTEC to CD4+SP | $9.99 \times 10^{-6}$ | $[1.00 \times 10^{-5}, 5.05 \times 10^{-4}]$ |
| $p_{mTEC}$ | Ratio of normal cells in the initial number of mTEC | $3.11 \times 10^{-4}$ | $[1.81 \times 10^{-5}, 6.91 \times 10^{-1}]$ |
| $\omega_{mTEC}$ | Death rate of irradiated mTEC | $1.62 \times 10^{-1}$ | $[3.10 \times 10^{-2}, 2.51 \times 10^{-1}]$ |
| $n_{mTEC}^{tot}(0)$ | Initial value for the number of mTEC | $1.44 \times 10^{5}$ | $[9.54 \times 10^{4}, 2.09 \times 10^{5}]$ |
| $\phi_{m}$ | Constant inflow to mTEC | $1.35 \times 10^{1}$ | $[1.23 \times 10^{1}, 5.29 \times 10^{2}]$ |
| $\phi_{m4}$ | Inflow to mTEC regulated by CD4+SP | $3.36 \times 10^{-4}$ | $[9.42 \times 10^{-5}, 3.31 \times 10^{-3}]$ |
| $r_{m}$ | Proliferation rate of mTEC | $5.39 \times 10^{-1}$ | $[7.17 \times 10^{-2}, 9.09 \times 10^{-1}]$ |
| $K_{m}$ | Carrying capacity of mTEC | $1.00 \times 10^{5}$ | $[2.66 \times 10^{4}, 1.09 \times 10^{6}]$ |

| | | | |
|---|---|---|---|
| $\gamma_{mp}$ | Negative regulation rate by DP to mTEC | $4.09 \times 10^{-9}$ | $[3.31 \times 10^{-9}, 1.96 \times 10^{-8}]$ |
| $\tau_m$ | Delay of DP regulating mTEC | $3.22 \times 10^{0}$ | $[2.56 \times 10^{0}, 4.92 \times 10^{0}]$ |
